# Convergence in sympatric swallowtail butterflies reveals ecological interactions as a key driver of worldwide trait diversification

**DOI:** 10.1101/2023.02.24.529892

**Authors:** Agathe Puissant, Ariane Chotard, Fabien Condamine, Violaine Llaurens

## Abstract

Ecological interactions may fuel phenotypic diversification in sympatric species. While competition can enhance trait divergence, other ecological interactions may promote convergence in sympatric species. Within butterflies, evolutionary convergences in wing color patterns have been reported between distantly-related species, especially in females from palatable species, where mimetic color patterns are promoted by predator communities shared with defended species living in sympatry. Wing color patterns are also often involved in species recognition in butterflies, and divergence in this trait has been reported in closely-related species living in sympatry as a result of reproductive character displacement. Here we investigate the effect of sympatry between species on the convergence vs. divergence of their wing color patterns in relation to phylogenetic distance, focusing on the iconic swallowtail butterflies (family Papilionidae). We developed a new unsupervised machine-learning-based method to estimate phenotypic distances among wing color patterns from 337 species allowing to finely quantify morphological diversity at global scale within and among species, allowing to compute pairwise phenotypic distances between sympatric and allopatric species pairs. We found that sympatry promoted strong convergence, while divergence in sympatry was less frequent and weaker. The effect of sympatry on convergence was stronger on females than males, suggesting that differential selective pressures acting on the two sexes drove sexual dimorphism. Our results highlight the significant effect of ecological interactions driven by predation pressures on trait diversification in Papilionidae and evidence the interaction between phylogenetic proximity and ecological interactions in sympatry acting on macroevolutionary patterns of phenotypic diversification.

## Introduction

Species interactions are an important driver of phenotypic diversification at both microevolutionary and macroevolutionary scale, promoting either convergence or divergence in sympatry. Antagonistic and mutualistic interactions indeed fuel the evolution of suites of traits involved in the adaptation to specialized ecological niches, therefore overcoming the effect of shared ancestry. Traits involved in interspecific interactions have often been shown to diverge in coexisting species, because of their partition into different niches (1,2). This effect has been extensively documented for traits involved in resource acquisition (3, 4). For example, in Plethodon salamander communities, cranial robustness and head shape divergence increased in sympatry compared to allopatry (5), probably in response to inter-specific competition. Phenotypic convergence can also be observed in sympatric species at large scales because of shared selective pressures. For example shared predator communities can favor convergence in warning coloration in Ithominii butterflies (6) and social interactions may promote song similarity in territorial birds (7).

Importantly, the effect of species interactions in the convergence vs. divergence of trait may depend on their level of relatedness. For instance, interspecific competition for territory (3) or mates (8) can promote either phenotypic convergence or divergence between sympatric species depending on their phylogenetic proximity. Closely-related species are indeed more likely to share similar ecological niches and to display similar suite of traits. Such resemblance may further increase interspecific sexual interactions. These reproductive interferences, from heterospecific courtship to mating producing unviable hybrids, are energetically-costly and thus favor reproductive character displacement in trait used as mating cues (9). Reproductive interferences may thus accelerate evolutionary divergence in traits involved in species recognition between closely-related species living in sympatry (10, 11). Interspecific interactions such as mimicry are thus more likely to trigger phenotypic convergence among phylogenetically-distant species. For example, flowers from species belonging to different families (*Turneraceae* and *Malvaceae*) may display similar coloration, because of increased attraction of local pollinators, without the cost of producing costly hybrids (12). The relative effect of ecological interactions (promoting convergence) and of phylogenetic proximity (enhancing the risk of costly heterospecific interactions) on overall pattern of phenotypic diversification is yet largely unknown. These opposite evolutionary forces generate an evolutionary trade-off (13), which may change the direction of phenotypic evolution in different species depending on the ecological effects generated by local communities of interacting species, and their level of relatedness. In diverse communities with different levels of relatedness in sympatry, is convergence generally promoted, or is divergence rather prevailing?

Within butterflies, convergence in wing color pattern can induce opposed effects between ecological interactions and phylogenetic distances. In unpalatable species, the evolution of mimetic color patterns in sympatry is promoted by predator learning and avoidance of prey harboring a known warning signal. This predator behavior generates positive density-dependent selection, favoring convergence of wing color patterns in sympatry, referred to as Müllerian mimicry (14, 15). The evolution of mimetic color patterns is also frequently observed in palatable species, referred to as the Batesian mimicry, where displaying a coloration close to those exhibited by defended prey provides a strong advantage (16). Both Müllerian and Batesian mimicries were shown to shape wing color pattern variations in butterflies, as for instance in the chemically-defended Heliconius species (17), or in the palatable species of the genus *Papilio* (18), respectively. Within *Papilio*, the powerful selection exerted by predators has promoted striking wing color pattern convergence among distantly-related species. For instance, striking resemblance can be observed between individuals from the unpalatable *Papilio polytes* and the chemically-defended *Pachliopta aristolochiae* (19), while these clades diverged about 50 million years ago.

Interestingly, butterfly wing color pattern is also involved in species recognition during mate choice, favoring their divergence between closely-related species living in sympatry (20). This trade-off between natural and sexual selection is likely to have different consequences on the evolution of the color pattern in the two sexes. For instance, Batesian mimicry is restricted to females in several species. The slower flight of egg-loaded females, as well as their predictable behavior of egg laying on specific host-plants may increase their predation risk, therefore further promoting Batesian mimicry in females (21, 22). Reproductive interference with other species living in sympatry may favor female preference for non-mimetic males, thus promoting mimicry in females and not in males (23). A recent study on sexual dimorphism in wing color patterns carried out on European butterflies indeed stressed out the major role of sexual selection in the evolution of color pattern in males (24). The evolutionary trade-off between convergence towards mimetic signal and divergence in mating cues might thus differ between sexes, and thus drive the evolution of sexual dimorphism. What evolutionary force prevails on the diversification of butterfly colour pattern? A global pattern of sympatric divergence is expected if sexual selection (or selection against sexual interference) prevails, while repeated convergences are predicted should natural selection imposed by predators play a predominant role.

Here we investigate the effect of sympatry on macroevolutionary pattern of color pattern diversification in swallowtail butterflies (Papilionidae), using 1,358 photographed individuals from 337 species distributed throughout the world. We specifically aim at testing the opposite effect of mimicry and reproductive interference and its interaction with phylogenetic proximity between interacting species. Wing color patterns displayed in butterflies are complex traits composed of various features, such as stripes, patches, rays of different shapes and colors, that may evolve in concert (25). To assess convergence and divergence in these complex color patterns, we develop a new machine-learning based method to quantify subtle variations in wing color pattern. This precise quantification of color pattern variations then allows comparing sympatric vs. allopatric species for both sexes, therefore testing for the relative effects of phylogenetic distances, ecological and sexual interactions on the evolution of color pattern in sympatry.

## Results

### 1. Quantifying color pattern variation using an unsupervised method

We quantified color pattern similarity within and among Papilionidae species without relying on any pre-existing human classification, through the development of an unsupervised similarity learning algorithm, based on the SimCLR method (a Simple Framework for Contrastive Learning of Visual Representations, 26). The similarity learning relies on modifications of the original images: the algorithm places the modified versions from the same image close to each other in the representation space, while modified versions from different images stand at larger distances. This unsupervised method allows to partly control the features used during classification: using image cropping and rotations to produce the modified images, we forced the model to ignore wing shape variations among species, while color pattern variations are used as discriminative features.

We used 2,716 standardized photographs of dorsal and ventral sides of Papilionidae butterflies and automatically separated the wings from the bodies. The model was then trained on the modified images of the four wings together standing on a neutral background. This training resulted in representation vectors of dimension 2,048 for each butterfly image, which contains information about the features displayed in the image. We then used a Principal Component Analysis (PCA) reducing the dimensionality to 20, while retaining approximately 80% of the variance. In certain species, several phenotypic forms are described, and we thus carried out the examination at the level of forms. We plotted the mean phenotype for each form within species in the resulting morpho-space (Figure 1 a,b). As there is substantial sexual dimorphism in Papilionidae, we first conducted independent analyses for males and females. As expected, interspecific distances were significantly much higher than intraspecific distances for both males and females, when studying either dorsal or ventral sides (Wilcoxon, p-value < 0.001 for males and p-value < 0.001 for females, Fig. S2), indicating that the model was able to successfully discriminate species based on phenotypic differences. Because dorsal and ventral pattern were very similar for most species, we then only show the analyses performed on dorsal wing color pattern. While our method does not allow to precisely identify the contribution of different wing features to the different PCaxes, the first axis clearly discriminates colored vs. white and black patterns. The gradient-based class activation mapping (Grad-CAM) for each picture then allows to pinpoint the pixels which were mostly used as discriminative features by the model (Figure 2). Elements of the pattern, such as stripes or color patches, show high activation, and were thus used as discrimination criteria by the model. In most species displaying hindwing tails, the tails were not considered as discriminant features (e.g. *Battus philenor, Graphium weiskei, Pachliopta aristolochiae, Papilio caiguabanus;* Figure 2 c, d, g, h). However, in some species where the hindwing tails are particularly prominent (e.g. *Lamproptera curius;* Figure 2 m), the model did rely on the tails as discriminant feature. Our phenotypic quantification thus mostly relied on color pattern variation rather than wing shape, but still accounted for large, visually-discriminant, wing shape variations.

**Figure 1.**
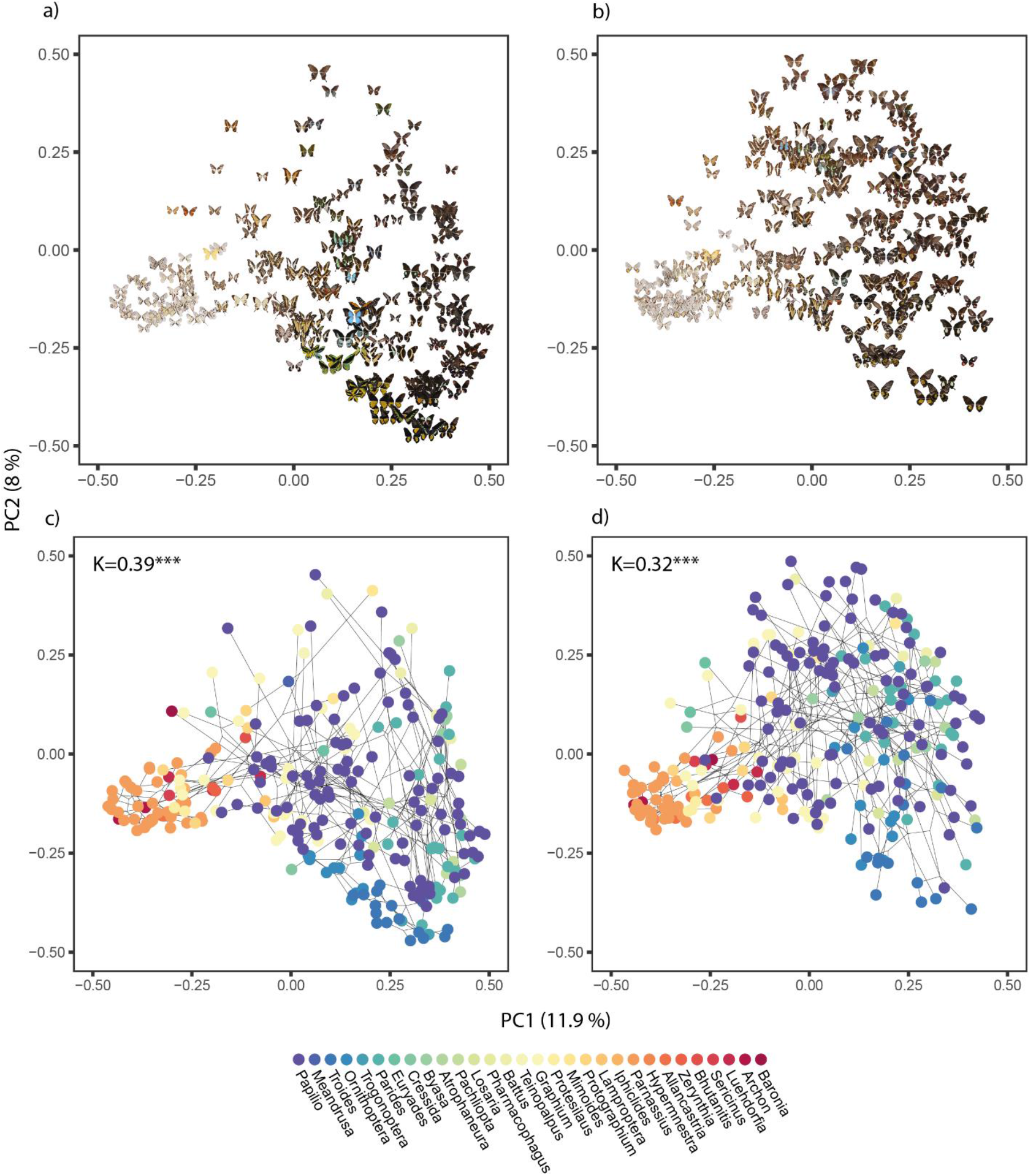
Color pattern variations and phylogenetic relationships across Papilionidae butterflies. First row: Phenotypic variations captured by the unsupervised machine-learning based methods applied to our 2716 pictures of Papilionidae. Independent PCA were carried out on (a) males and (b) females. Note that we used the mean phenotype by sex and by species and represented only the first two axes, explaining only 19.9 % of the phenotypic variance. We display the actual picture of butterfly from each form and each sex on the morphospace to observe how actual color pattern variation was separated by our method of phenotypic discrimination. Second row: Phylo-morpho space computed on the mean phenotype for each species in (c) males and (d) females. The phylo-morpho space is the projection of the morpho space coordinates on the first two axis of the morpho space principal components. Each colored dot represents the phenotype of a given species and the black lines show the projection of the phylogenetic relationships among the species. The color code corresponds to the different genera, the color gradient corresponding to the location of the genus on the phylogeny.

**Figure 2.**
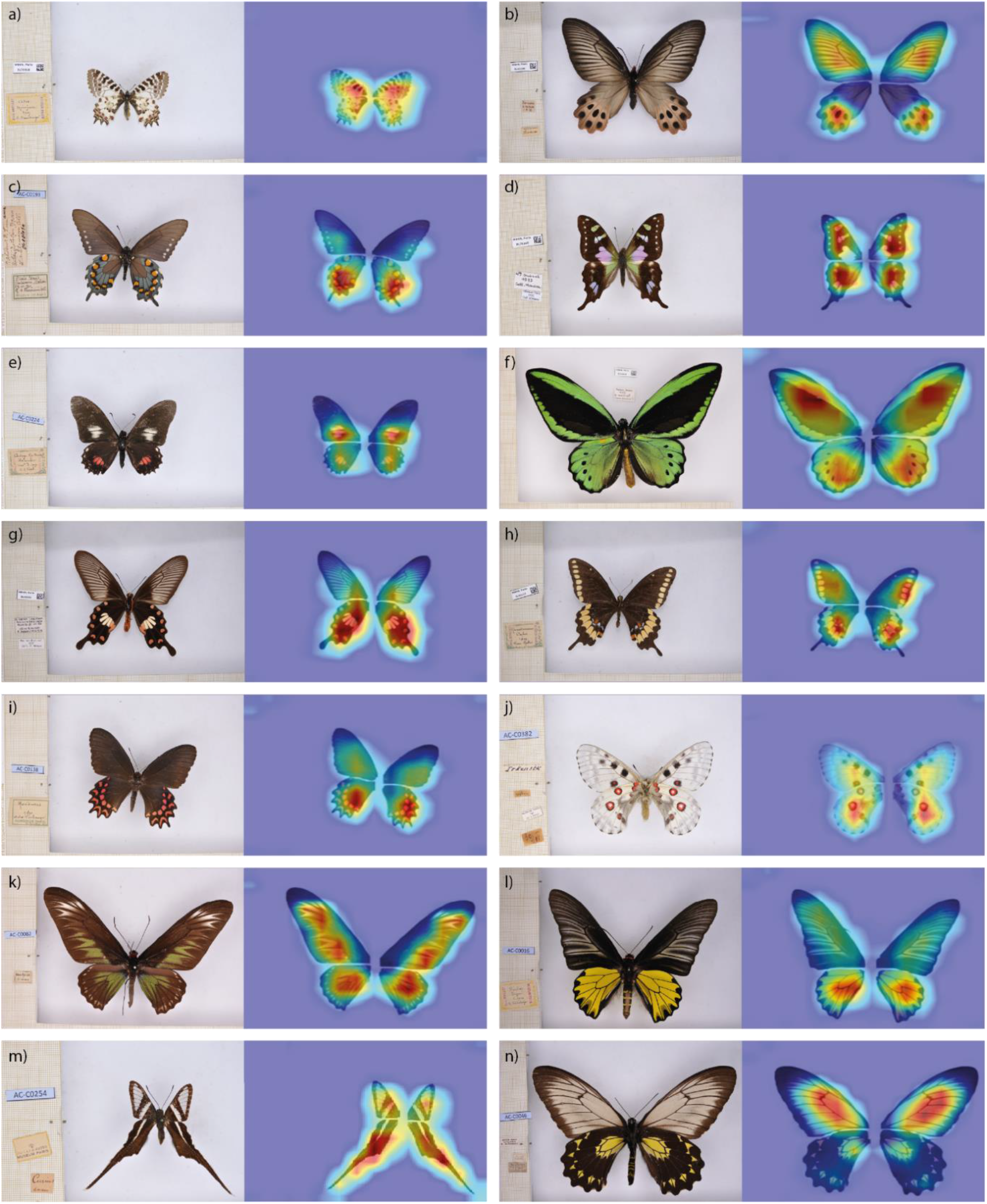
Discriminant features used to build the morpho-space obtained using Grad-CAM mapping. We selected one or two species for several genera to give examples that cover different parts of the phenotypic space. Redder color indicates pixels that weights the most in the activation of the neural network. From left to right, up to down: Allancastria cerisyi, Atrophaneura priapus, Battus philenor, Graphium weiskei, Mimoides euryleon, Ornithoptera priamus, Pachliopta aristolochiae, Papilio caiguabanus, Parides photinus, Parnassius nomion, Trogonoptera brookiana, Troides aeacus, Lamproptera curius, Troides andromache.

### 2. Phylogenetic and geographic patterns of phenotypic disparity

To test for the effect of phylogenetic proximity on color pattern divergence, we then built a phylo-morphospace for each sex (Figure 1 c,d). These morphospaces show the position of the average dorsal color pattern within each species, and a projection of the phylogenetic relationships among species using the principle of unscaled squared change parsimony, based on the most recent and complete swallowtail phylogeny (27). We detected a significant phylogenetic signal on phenotypic variations in both males (K = 0.39, p-value < 0.001) and females (K = 0.32, p-value < 0.001). K values lower than 1 for both sexes indicate that closely-related species were generally less phenotypically-similar than expected under a Brownian motion model of phenotypic evolution. The phylogenetic signal was also lower in females than in males, suggesting that female phenotypes tend to diverge more from the Brownian expectation than male phenotypes, pointing at differences in the evolutionary forces shaping phenotypic evolution between the two sexes.

To test for spatial variation in color pattern disparity, computed as the average squared distance of phenotypes from the centroid of phenotypes, we then gathered the geographical range distribution at the species level for 225 out of the 267 species where both male and female phenotypes were sampled. By mapping the phenotypic disparity on the world map and comparing it with species richness, phylogenetic diversity (Fig. S5), we observed that the areas of highest disparity were in Europe and Middle East. The spatial variation in disparity therefore did not match the documented hotspots of species richness: the correlation between spatial pattern of disparity and local phylogenetic diversity was indeed poor (Spearman rank correlation on males: 0.48, females: 0.49).

The mimicry between species, and especially female-limited mimicry documented in Papilionidae, is likely to account for the limited phylogenetic signal in color pattern evolution and the limited impact of species richness on local disparity. To test the effect of mimicry between species on macroevolutionary pattern of wing color pattern diversification, we thus investigated the effect of sympatry between species on color pattern evolution.

### 3. Convergence events are more frequent in sympatry

To test for the effect of sympatry on phenotypic evolution, we defined species that shared parts (≥20%) of their geographical range as sympatric, while all other species with non-overlapping distribution (<20%) were classified as allopatric. We then designed a permutation test to detect phenotypic convergence and divergence.

We first established a null distribution by permuting the residuals of the regression between the phenotypic and phylogenetic distances, throughout each pair of phenotypic form within and among species. We then tested for each pair of forms whether they were more or less similar than expected from the null distribution, corresponding to convergence and divergence, respectively. We quantified the strength of convergence and divergence from the difference between observations and the generated null distribution. We tested each pair for phenotypic convergence and divergence irrespectively of any other factors, such as geographical range distribution.

In males, out of the 29,402 pairs, we detected 256 convergent pairs and 193 divergent pairs, including 25 and 7 in sympatric pairs, respectively. In females, out of the 31,877 pairs, we detected 281 convergent pairs and 220 divergent pairs, including 55 and 7 in sympatric pairs, respectively. The relative number of convergence vs. divergence events detected was significantly larger in sympatric pairs as compared as in allopatric pairs, for both males and females (χ^2^= 4.54, p-value < 0.05 and χ2= 25.37, p-value < 0.001, respectively).

Moreover, we found that convergent pairs in sympatry were in general more distantly related (on average 93.96 Myr – sum of branch lengths – for males and 84.54 Myr for females) than divergent pairs in sympatry (in average 41.61 Myr for males and 42.54 Myr for females; Wilcoxon, p-value < 0.05 for females and p-value < 0.01 for males). Because of our phylogenetic correction, convergence is indeed more likely to be detected in distantly-related pairs, because their phenotypic resemblance cannot be explained by shared ancestry. Conversely, divergence is less likely to be detected in these distantly-related species. To disentangle the effect of the phylogenetic correction from the actual effect of mimicry on convergence, we then investigated the strength of convergence in males and females.

### 4. Evolution of sexual dimorphism driven by the evolution of female color pattern

Focusing on detected convergence events, we did not find significant differences in convergence strength between sympatric and allopatric species pairs for males. In females, however, the convergence was significantly stronger in sympatric species pairs (Wilcoxon, W = 7448, p-value < 0.05). Consistent with the lower phylogenetic signal of color pattern in females as compared to males, our test of convergence suggests that ecological interactions between sympatric species have a more intense effect of on the evolution of female vs. male phenotypes.

To investigate the evolutionary divergence between male and female phenotypes within species, we estimated sexual dimorphism by computing the Euclidean distances in the PCA space between the two sexes within each species. We also computed the raw contrasts for each pair of sister species – male/male distance and female/female distance in sister species – and studied the ratio of male contrast over female contrast (r): when the ratio r < 1, female phenotypes diverged more than male ones in the same amount of time (Figure 4). We investigated the distribution of relative divergence of color pattern in each sex in dimorphic species to test for the relative effect of selection generated by reproductive interference on male phenotypes vs. ecological selection in female phenotypes on the evolution of sexual dimorphism. Dimorphic species were defined as species with dimorphism value above 0.3. We found that most of the dimorphic sister-species pairs indeed had r < 1 (one sample Wilcoxon test, V = 621, p-value < 0.05), indicating that the sexual dimorphism in color pattern mostly stems from the divergence of female phenotype. However, the distribution shows high values for a limited number of observations: in a few rare cases, the dimorphism was driven by the divergence of male phenotypes. When sexual dimorphism was driven by the evolution of male phenotype, the level of phenotypic divergence away from the ancestral color pattern was much higher than when the sexual dimorphism was driven by the divergence of female phenotype. This confirms that female phenotypes generally diverge more than male ones resulting in sexual dimorphism driven by ecological selection acting on females. Nevertheless, sexual dimorphism could still arise from reproductive interference fueling sexual selection promoting male phenotypes differing from heterospecific males and generating very strong divergence in male phenotypes.

**Figure 3.**
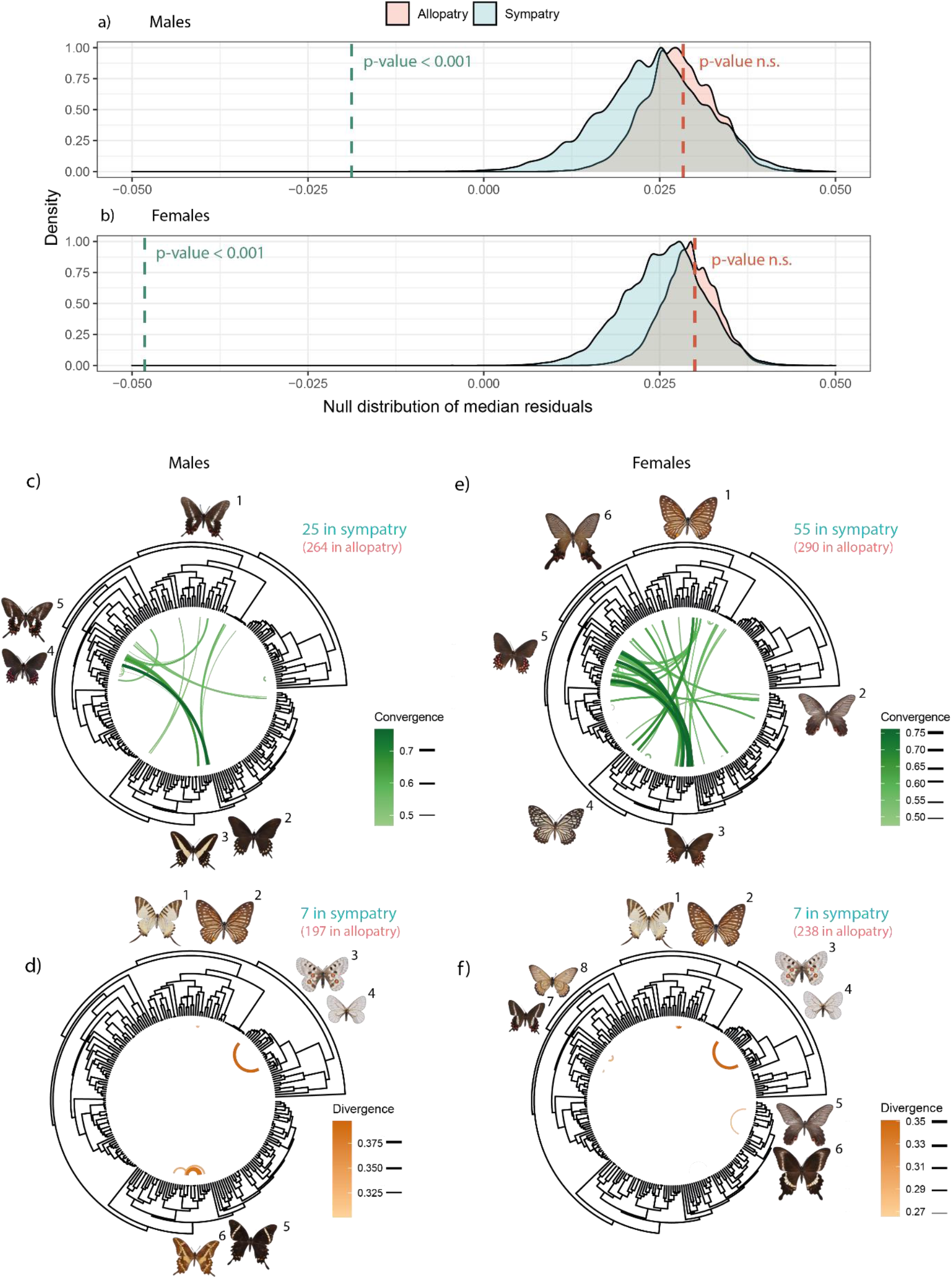
Detection of convergence and divergence events between sympatric and allopatric species pairs. Distribution of residual medians for (a) males and (b) females after 100,000 permutations for sympatric pairs in blue and allopatric pairs in pink. The dashed blue vertical line corresponds to the observed median for the residuals of pairs of sympatric species, and the dashed pink vertical line corresponds to the observed median for the pairs of allopatric species. (c-f) Phenotypic convergence and divergence associations between pairs represented on the swallowtail phylogenetic tree with male convergence (c) and divergence (d) between pairs, and female convergence (e) and divergence (f) between pairs. Example of pairs of convergent and divergent species are plotted along the phylogeny: c-1: Mimoides lysithous, c-2: Papilio erostratus, c-3: Papilio hectorides, c-4: Parides photinus, c-5: Parides bunichus. d-1: Graphium antiphates, d-2: Graphium xenocles, d-3: Parnassius nomion, d-4: Parnassius stubbendorfii, d-5: Papilio pelaus, d-6: Papilio aristodemus. e-1: Graphium xenocles, e-2: Papilio protenor, e-3: Papilio erostratus, e-4: Papilio clytia, e-5: Parides photinus. f-1: Graphium antiphates, f-2: Graphium xenocles, f-3: Parnassius nomion, f-4: Parnassius stubbendorfii, f-5: Papilio protenor, f-6: Papilio hipponous, f-7: Parides bunichus, f-8: Euryades corethrus.

**Figure 4.**
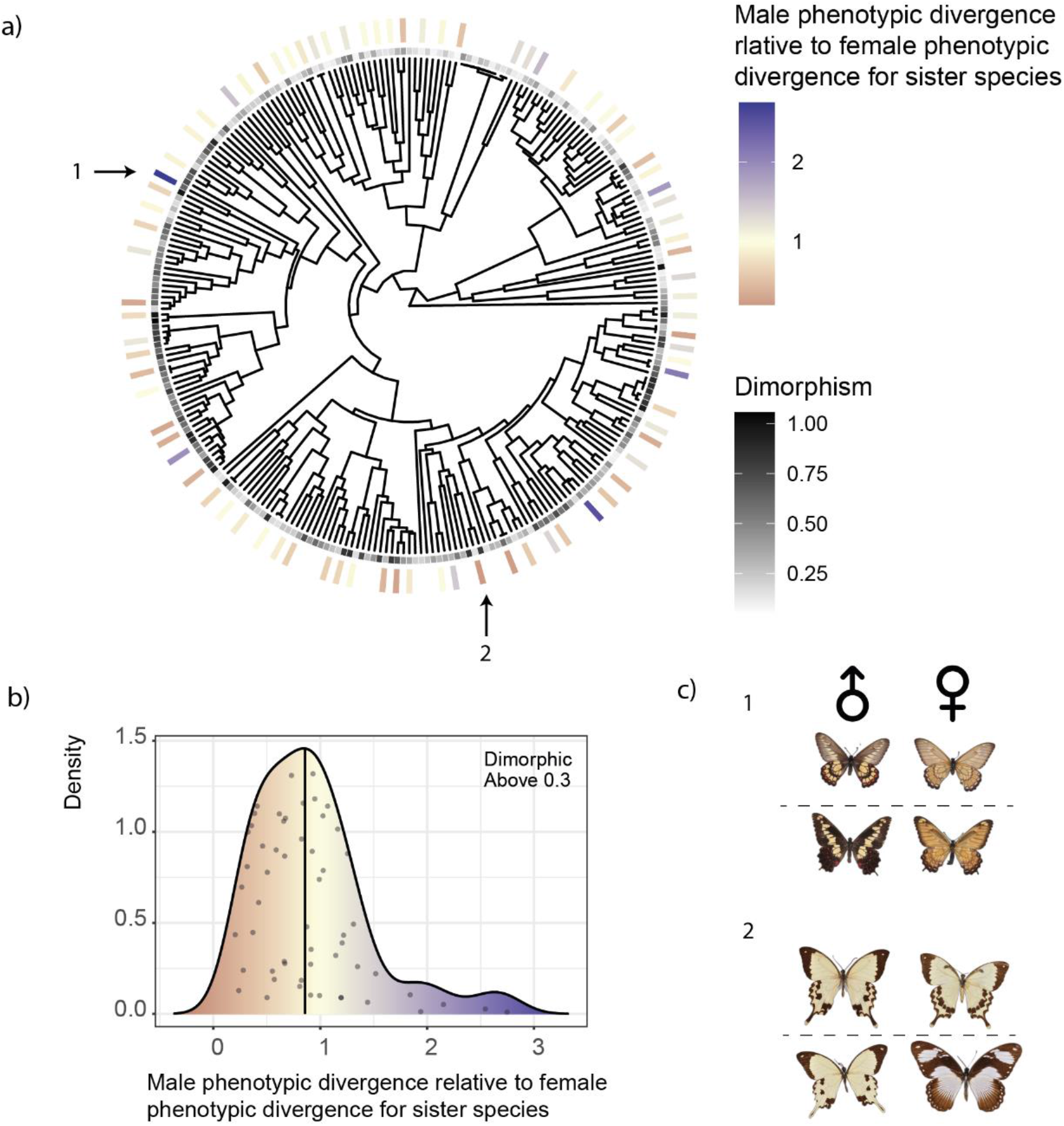
Evolution of sexual dimorphism in color pattern throughout the Papilionidae family: contribution of divergence of male phenotype vs. female phenotype in the evolution of sexual dimorphism was estimated by comparing sister-species. a) Swallowtail phylogeny showing sister-species with the level of dimorphism (grey scale) and ratio of raw contrasts: male/male distance over female/female distance for sister species (gradient from red indicating a small ratio value to yellow indicating a ratio of 1 to blue indicating big ratio value). b) Distribution of the ratio of raw contrasts for sister species with the median indicated. c) Male and female specimens are shown for the two-sister species presenting the highest ratio: (1) Euryades corethrus above and Euryades duponchelii below, and for the two sister species presenting the lowest ratio: (2) Papilio meriones above and Papilio dardanus below.

## Discussion

Using a novel machine learning based method, we quantified color pattern variation at the global geographic scale in Papilionidae butterflies and uncovered a general trend of convergence among sympatric species. This method is not specific to butterflies and could be trained to quantify variation in visual signal in any biological model. By tuning the modifications made during the training, the method can be trained to quantify shape, color, and pattern similarity in various organisms, opening new research avenues for investigation phenotypic diversification.

### 1. Uncovering discriminant features of visual signal

In this study, we provide the first quantitative analysis of butterfly color pattern variation at a large phylogenetic scale through the comparison of highly divergent and complex color patterns. The complexity and extreme diversity of butterfly color patterns has led previous studies to simplify the problem. Some authors have discretized the patterns (21,28), focusing on a few features at a time (29, 30). This approach comes at the cost of losing part of the signal as well as its global integration. Other approaches that rely on the alignment of wing images on the color patterns themselves allow powerful comparisons based of whole color patterns (31). The alignment is however limited to closely similar patterns and wing shapes, thereby preventing large evolutionary scale analyses. Here we circumvent these issues with a new machine learning method to automatically quantify color pattern variation among distantly-related species in an unsupervised way, independently from any pre-existing human classifications.

This unsupervised and automatic method allows comparing multiple color patterns among very distantly-related species, which may differ in wing shape because of other selective constraints - see Le Roy et. al. (2019) (32) for a review. We thus designed our method to neglect minor wing shape variation (see Sup. Mat Figure 7 showing the limited effect of wing shape variations on our estimates of distances between color patterns). Nevertheless, wing shape variation is not independent from color pattern and does contribute to visual signal perceived by congeners and predators. Linke et al. (2022) (33) showed that wild blue tits can learn to associate both color pattern and hindwing tails with escape ability, highlighting the relevance of prominent shape features such as wing tails in the generalization performed by visual observers. Although our method reduces the impact of minor wing shape variation, it still retains information about prominent discriminative parts of the wings such as large tails (Fig. 2). Our method thus provides relevant quantification of color pattern variations within and among species for studying how selection exerted by visual predator and/or congeners shapes the diversification of wing color patterns.

Nevertheless, we quantified visual signal similarity in the visible light range only. Both butterflies and some of their avian predators are UV-sensitive, and some patches found on Papilionidae wing patterns do reflect UV wavelength (e.g. in the white patches of Papilio glaucus, 34). Pigments or structural changes in wing scale that lead to UV reflection generally also generate differences in the visible reflectance, but variation in the UV reflectance may lead to conspicuous differences for UV-sensitive species, that cannot be captured by our method. However, color pattern mimicry often involve convergence in the UV pattern as well (as shown in Papilio polytes, 35): ignoring the UV signal would thus not strongly modify the broad trend of wing pattern convergence in Papilionidae.

### 2. Phylogenetic distances interfere with the ecological interactions

As we detected weak but significant phylogenetic signal on wing color variations, we used a linear regression to control for the effect of phylogenetic distances in our analyses. We found that convergent pairs were more distantly related compared to divergent pairs, and that convergence strength was higher than divergence strength. Nevertheless, this phylogenetic correction might bias our detection of convergence and divergence events. When closely-related species display a strong phenotypic similarity, it is challenging to disentangle the effect of shared ancestry from selection promoting color pattern convergence. Conversely, when distantly-related species display very different phenotypes, the relative effects of phylogenetic distance from divergence due to selection are confounding. The necessary phylogenetic correction therefore limits the detection of (1) convergence events among closely-related species, and (2) divergence among distantly-related species. The greater number of convergence events detected in our study among distantly-vs. closely-related species might stem from decreased reproductive interferences in phylogenetically distant mimetic species but might also stem from the bias induced by the phylogenetic correction. Nevertheless, the significantly greater strength of convergence as compared to divergence in sympatry estimated from our analyses despite phylogenetic correction suggests that natural selection promoting convergence in sympatric species is a significant evolutionary force interacting with neutral divergence. Other measurement of convergence, such as the C1 statistic (36), are also a distance based statistic with permutations to assess significance have the same phylogenetic bias. This C1 statistics was recently used to assess convergence among models and mimics within communities (37) but is more computationally intensive than our method. When computing the C1 statistic for our convergent pairs and comparing our convergent strength, we found similar results.

Interestingly, we found that sympatric pairs tended to be more closely related than allopatric pairs (Wilcoxon, males: p-value < 0.001, females: p-value < 0.001). Recently diverged species indeed tend to retain similar habitat affinities (phylogenetic niche conservatism) and species from a same clade thus frequently occupy similar geographic regions (38). We found convergence in sympatric pairs despite this phylogenetic clustering. While we cannot directly assess whether trait convergence is due to shared environmental conditions versus species interactions, the phylogenetic correction should account for trait similarities due to the combination of phylogenetic constraints or phylogenetic niche conservatism (39). The detected convergence is thus likely to stem from local selection exerted by predator behavior.

### 3. The effect of sympatry on phenotypic convergence and divergence

Negative interactions in sympatry such as competition for resources and territory are often assumed as the main driver of phenotypic evolution, in line with character displacement theory (1, 2). Contrastingly, we found an effect of sympatry on trait convergence among species. Overall, among all the detected convergent pairs of sympatric species, 60% were previously mentioned in the literature as potential mimetic species (see the detailed list of all convergent and divergent sympatric pairs, and corresponding references in supplementary materials). Despite our simplistic definition of sympatry, based on geographic range overlap without precise information on the shared micro-habitat, our method conducted at the global scale did manage to recover large number of species pairs identified as mimetic in field studies. Female-limited Batesian mimicry has indeed long been documented in Papilionidae (19, 40), and our study recovers well-studied cases, such as the documented Batesian mimicry between Amazonian unpalatable *Parides* species, and their *Papilio* mimics – see for example the black pattern with conspicuous red patches in the unpalatable *Parides photinus* and its mimic *Papilio erostratus* (18) (Tab. S2). Interestingly, convergence was also detected within the *Parides* genus, for example between *Parides photinus* and *Parides montezuma*, which reinforces the hypothesis of a Müllerian mimicry ring within the *Parides* genus (41). Convergence was also found between Southeast Asian species *Graphium xenocles* and one form of *Papilio clytia*, both displaying a white color with contrasting black venation and an orange dot on the hindwing. While it is probable that both species are palatable, they are both mentioned as being mimics of unpalatable Danaine species and may thus belong to a same mimicry ring. Our study also suggests new convergence events, especially in females of sympatric species pairs. For example, we found convergence between the female flava form of *Papilio deiphobus* and the female of the unpalatable species *Troides rhadamantus*, both observed in the tropical broadleaf forests of Philippines. *Papilio deiphobus* is closely related to the species *Papilio memnon*. In *P. memnon*, female-limited polymorphic mimicry with different *Troides* species has been suggested (40). Our study thus points a potentially new events of convergence between the clade *Papilio* and *Troides*. Surprisingly, we also detected convergence in males from sympatric species pairs. First, there are convergence between males from species where convergence in females were also detected, for example in *Parides photinus* and *Papilio erostratus* described above. The convergence in male phenotypes may stem either from Batesian mimicry, but it may also stem from correlated color pattern evolution between males and females within the same species, caused by developmental constraints. In some species, convergence was detected among palatable species, without any evidence for mimicry with defended species. Instead, these convergent species exhibit color patterns composed of contrasting black rays on a clear background and orange and blue patches on the hindwing close to the tails, such as in the convergent *Papilio alexanor* and *Iphiclides podalirius*. This kind of color patterns combined with hindwing tails may contribute to the deflection of predators away from the vital parts of the body (42). Such convergence in color pattern could thus be driven by predator cognitive bias and associated behavior but do not imply mutualistic or parasitic interactions between butterfly species.

Such a strong effect of trait convergence in sympatry at large evolutionary scale was reported for the songs produced by ovenbirds (family Furnariidae), most probably explained by ecological interactions between sympatric species (7). On the contrary, sympatry did not have any significant effect on the evolution of several traits in haemulid fishes, including their color patterns (43). Interestingly, a study focusing on color pattern variations in sympatric species of newt revealed convergence in dorsal patterns, involved in concealment from predators and divergence in the ventral colors, likely in relation with mate choice (8). In Papilionidae, the dorsal wing color pattern is likely involved in interactions with both predators and mating partners. However, the global increase in convergence detected in sympatry suggests that selective pressures linked to predation are prevalent over reproductive character displacement in driving color pattern evolution, especially in females.

### 4. Sex-specific selection pressures and the evolution of sexual dimorphism

Assuming that heterospecific mating is costlier for females than for males, mimicry promoted by predator behavior is more likely to emerge in females than in males (23). In Papilionidae, we found that significantly more and stronger cases of convergence are detected in sympatry compared to allopatry for females, but not for males. The phylogenetic signal associated with color pattern variation is higher in males than females. This is consistent with the previously described high prevalence of female-limited Batesian mimicry in Papilionidae. For instance, in the genus *Papilio*, approximately 25% of species have mimetic females, while males generally display the ancestral color pattern (44). We found that overall, female color patterns diverged more than male color patterns in sister species. The color pattern dimorphism then seems to be mostly driven by natural selection acting on female phenotypes (21) but may also be favored by female preferences towards males displaying ancestral phenotypes, limiting heterospecific interactions.

In the rare cases where male color pattern diverged more than female ones in sister species, the divergence of male phenotype away from the ancestral color pattern was very strong, for example in the sister species *Euryades corethrus* and *E. duponchelii* (see Figure 4). Such a strong divergence in male ornament could be fueled by runaway process (as described by Fisher in 1930, 45). When a given trait displayed by males is preferred by females, this leads to an increased number of offspring carrying both the preference and the preferred color pattern, and to an acceleration in the evolution of the male trait (46, 47). The evolution of male traits and female mate preference evolution reinforce one another, leading to the evolution of “extreme” ornaments (48). The high divergence in male color pattern away from the ancestral pattern and the phenotypes of female observed in some species, might thus be driven by sexual selection by females provoking an acceleration in the evolution of male phenotypes, away from the phenotypes displayed in males from closely-related species living in sympatry.

Our method is nevertheless likely to fail for detecting some cases of sexual dimorphism driven from divergence in male phenotypes in sympatric species, implying UV reflectance: some sister species with partially overlapping geographic distribution, like *Iphiclides podalirius* and *I. feisthamelii*, have highly similar color patterns but males differ in the UV reflectance of some patches, while no such differences were detected in females (49). This divergence in male coloration might stem from selection generated by reproductive interference in these parapatric sister-species. Our method aims at detecting sexual dimorphism generated by ecological interactions in sister-species. This focus on sister-species discards the effect of the phylogenetic correction bias but prevents the detection of ancestral sexual dimorphism. For example, the strikingly sexual dimorphism observed in the *Ornithoptera* genus, where males display brightly saturated green, blue and orange colors and females are more melanic and cryptic, is not detected in our test as stemming from male divergence. Such ancestral sexual dimorphism that might stem from sexual selection acting on males is likely to be independent from reproductive interference between sympatric species.

### 5. Conclusion

The method described here allowed to uncover global trend at a large phylogenetic scale, and is likely to stimulate new studies on the diversification of visual traits that were challenging to compare so far. Our study on the diversification of color pattern in Papilionidae indeed showcases the major role of ecological interactions among sympatric species in trait diversification and shed light on the impact of phylogenetic distances on these interactions, leading to contrasted patterns of convergence and divergence in sympatry.

## Materials and Methods

### 1. Sampling and standardized photographs

We sampled specimens from the collection of the National Museum of Natural History (Paris) to cover most of the described Papilionidae species. We relied on the latest taxonomic reviews (50) and published dated species-level phylogeny (27) to scan the collections for the described species. We sampled 337 species out of the 382 shown in the phylogeny of Allio et al. (2021) (27). For each species, we selected two males and two females, whenever specimens were available. 17 species presented several phenotypic forms (2 in average for a total of 32 forms, which we sampled). Our sample was thus composed of 1,358 individuals, including 774 males (329 species) and 592 females (273 species; note that females are much rarely collected so that female were lacking for many species). The dorsal and ventral side of the sampled individuals were photographed under controlled LED light using a Nikon D90 (Camera lens: AF-S Micro Nikkor 60 mm 1:2.8G ED), in standardized conditions. For the statistical analysis, we kept only species for which we have at least one male and one female, leading to a final species sample size of 267 species, 292 subspecies, and 296 unique phenotypic forms for males and 313 for females.

### 2. Wing segmentation

On each picture, the four wings were first digitally separated from the body and from the background of the pictures using a combination of machine-learning-based segmentation and traditional image processing. The machine-learning-based segmentation was performed using a Mask-RCNN model, which learns to classify each pixel on the picture either as belonging to the region of interest or not and generates a segmentation mask based on this classification. First, a training database of 371 pictures was constituted, by manually cropping the four wings on each picture. A Mask-RCNN model was trained on two thirds of this database (247 pictures) using the PixelLib python library. The rest of the pictures were used to evaluate the segmentation using the intersection over union of masks metric (IOU) with a threshold of 0.9 to consider the prediction a true positive. The IOU with this threshold of 0.9 was 0.85, *i.e*. for 85% of the predicted masks, the mask was 90% or more in agreement with manual cropping. As the produced masks often kept a few background pixels at the edges of the wings, a supplementary step was added, using traditional image processing. After this post-processing step, the IOU with a threshold of 0.9 went up to 0.98, meaning that 98% of the predicted masks were consistent with 90% of the manually cropped images. However, in a few pictures, wing parts with coloration closely matching the background color had to be manually eliminated. Finally, we obtained the masks for the four wings for each of the 2,716 standardizes pictures, and generated images with a single wing on a white background.

### 3. SimCLR training and evaluation

#### The SimCLR method

The similarity of color patterns was quantified using the new unsupervised deep contrastive metric learning method SimCLR (26). This method is designed to provide a distance metric between images and can produce the images’ coordinates in a space with reduced dimension. Traditionally, supervised contrastive learning needs labels for the images to determine whether pairs of images are considered similar (positive pair, belonging to the same class) or dissimilar (negative pair, belonging to different classes), allowing the neural network to learn features making images similar or dissimilar via modification of the neurons weight during training to obtain a smaller distance for positive pair and a larger one for negative pairs. SimCLR does not require such labelling of the pairs but considers each image as their own class, using modified versions of the image to perform comparisons. These modifications, referred to as image augmentations, include cropping, rotations, inversions, and color jittering, happening at a rate set by the user. Augmentations of the same initial image are considered as positive pairs, while augmentations of different images are considered as negative pairs. The metric learning is computed by minimizing a loss function (Normalized Temperature Cross Entropy Loss or NT-Xent loss). The loss is minimized by tuning the weights of the neural network during learning to increase the cosine similarity within positive pairs and to lower it in negative pairs. The cosine similarity between two vectors *u* and *v* is defined as 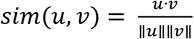. For a positive pair (*i,j*) in a batch of size *N*, the NT-Xent loss is defined as:

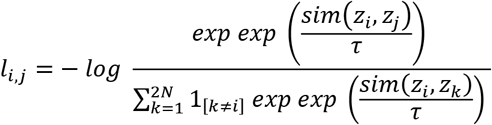

where 1_*k*≠*i*_ ∈ {0,1} is an indicator function evaluating to 1 if *k* ≠ *i* and *τ* denotes a temperature parameter.

#### Image augmentations

The choice of augmentations is crucial as it determines the features to which the method will be invariant. To study variation in color pattern, we thus applied randomized cropping with a minimum size of 50% of the original image (invariance to size); rotation, horizontal and vertical flipping each with a probability of 50% (invariance to left/right/up/down or orientation); conversion to grayscale with a probability of 20% (invariance to pattern without accounting for color); and finally, no color jittering of any kind to account for the color hue, brightness, and saturation of the actual wings.

#### Network’s architecture

Based on Chen et al. (2020) (26), we chose a ResNet50 (residual neural network with 50 convolutional layers) as the backbone of the network and then replaced the classification head with a SimCLR multilinear projection head. The Resnet50 backbone was initialized as pre-trained on the large ImageNet database to increase performance and compensate for the limited number of images in our own dataset. Three hyperparameters were optimized: the batch size, the number of training epochs and the temperature parameter. A grid search was performed to obtain the optimal parameter values, e.g., the parameters leading to the best performance during evaluation.

#### Evaluation & training

After the training phase, the output of the final convolutional layer was used as the vector representation of the images, as it retains more information than the output of the multilinear projection head. The evaluation relied on a pretext downstream task aiming at classifying the vector representation of images into different categories. To obtain categories from our images without *a priori*, we classified our images into 16 clusters, using the HDBSCAN clustering method (51) on vector representation obtained with a classical pre-trained convolutional neural network, VGG16 (52). The cluster labels were then used as class labels during classification. The backbone of the SimCLR method was also pre-trained on the same dataset. Contrary to the SimCLR method, the vector representations from the VGG16 network did not provide a relative metric space. Finally, to assess if the pre-training biases the evaluation, we compared performance of the pre-trained-only method, the pre-trained and finetuned during classification method, and the SimCLR trained method. A more in-detail discussion about the evaluation of the method can be found in supplementary materials. The method was implemented in Python, mainly using the Pytorch library for machine learning and using the Lightly library for SimCLR related augmentations, backbone, and loss function. Finally, the method was trained with a batch size of 128, several training epochs of 300 and a temperature of 0.5, obtaining a f1 score 1.15% higher than the pre-trained-only model, a kappa score 1.19% higher, and a mean accuracy of 95% (84% for the pre-trained-only method). A PCA then allowed to reduce the dimensions of the representation vectors from 2,048 to 20, while retaining approximately 80% of the variance.

### 4. Explainability and quality control

To identify the features of the images used in discrimination by the SimCLR method, we generated a gradient-weighted class activation mapping (Grad-CAM, 53) for the input images. This allows to pinpoint pixels of the image generating the highest activation in the convolutional layers. To check for the reliability of the image embeddings, the distances between all pairs of images were calculated. We then compared the distribution of intra-specific and interspecific pairwise distances to test whether images from butterflies belonging the same species had a lower phenotypic distance than interspecific pairs of images.

### 5. Species geographical range

For 125 out of the 267 species studied here, their geographical range was retrieved from the IUCN Red List Spatial Data & Mapping Resources (54) or from the Map Of Life project (55) from various datasets. When the geographical range could not be retrieved from those sources because they were missing or incomplete (101 species) the geographical range was estimated from GBIF occurrences by generating a convex alpha hull with a buffer distance around the occurrences. GBIF occurrence data were downloaded and cleaned using the rgbif and CoordinateCleaner R packages (56) and species with less than 30 occurrences were discarded. For 45 species, geographical range could not be retrieved because of missing or too few GBIF data. Those species were discarded from the analysis. The pairwise species overlap was calculated using the Jaccard index (area of the intersection of the ranges over area of the union of ranges). A pair of species was considered sympatric if their ranges overlap was 20% or more, and if not, it was considered allopatric.

### 6. Detection and quantification of convergence and divergence

#### Convergent and divergent pairs

To detect and quantify convergence and divergence, we designed a permutation-based method similar to the one described in (57). Because many species display multiple forms, we computed mean form phenotypes as the mean of the vectors of all specimens belonging to the same form. To assess the level of phenotypic similarity independently from phylogenetic proximity, we performed a linear regression between pairwise phenotypic distances (calculated as the Euclidean distance between the phenotypes) and pairwise phylogenetic distances. Negative residuals represent pairs for which phenotypic distance is lower than expected given the phylogenetic distance between species, which may indicate putative phenotypic convergence. Conversely, positive residuals indicate a larger phenotypic divergence than expected from phylogenetic distance. To identify the level of wing color pattern convergence in the swallowtail phylogeny, we tested whether the studied pairs were more phenotypically similar than expected at random, while controlling for phylogenetic distances. We thus compared the residuals obtained above to a null distribution of residuals. The null distribution was generated by permuting the residuals associated with each species throughout the phylogeny. The permutations were performed using the Lapointe-Garland permutation method, in which pairs of trait values are exchanged with probability inversely proportional to their phylogenetic distance (58). This permutation methods allows to correct for non-independent and identically distributed data, therefore removing phylogenetically induced false positives and accounting for unbalanced phenotypic distribution. Residuals were permuted 100,000 times over the 29,402 possible pairs for males and 31,877 for females. The *p*-value associated with phenotypic convergence for each pair was calculated as the proportion of permutations where the observed residuals was lower than the permuted residuals. We fixed the *p*-value at 1% and thus deemed convergence between a pair of phenotypic forms significant if 99% of the permuted residuals for this pair were higher than the observed residual. Conversely, to test for phenotypic divergence events, we performed the same permutation test but computed for each pair the proportion of permutations where the observed residual was greater than the permuted residual to obtain a *p*-value and fixed the *p*-value at 1%. Convergence and divergence strength were then quantified as the departure of the observed residuals from the median computed in the generated null distribution.

#### Sympatry and allopatry

To assess the overall convergence or divergence of sympatric pairs, we computed the median of sympatric pairs’ permuted residuals for each permutation. A *p*-value was computed by counting the number of permutations where the permuted median for sympatric pairs was higher than the observed median for sympatric pairs. To compare sympatry with allopatry, the same was done for allopatric pairs.

#### Dimorphism in sister species

To assess which sex phenotype drives the evolution of dimorphism, we determined which sex diverged phenotypically more than the other in the same amount of time for each pair of sister species. We computed the Euclidean distance between male phenotypic coordinates and the Euclidean distance between female phenotypic coordinates from the two species of each pair, which correspond to the raw contrasts for males and females respectively. To compare the two, we computed the ratio 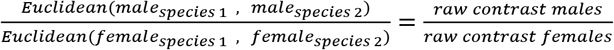 for each sister species.

When the ratio is close to one, males and females diverged equally in the same amount of time. When above one, males diverged more than females, driving dimorphism in the pair. When less than one, females diverged more than males in the same amount of time, driving dimorphism in the pair. We assessed the level of dimorphism in each species by taking the Euclidean distance between males’ phenotypic coordinates and females phenotypic coordinates.

## Data availability

All the code necessary for the python machine learning training and the R analysis, as well as the phenotypic coordinates in the morpho space are provided in the following Github: https://github.com/AgathePuissant/SimCLR_phenotypic_convergence

## Acknowledgments

This work was funded by UMR 7205 from Muséum National d’Histoire Naturelle. We thank the intensive computing platform « Plateforme de Calcul Intensif et Algorithmique PCIA, *Muséum national d’histoire naturelle, Centre national de la recherche scientifique, UAR 2700 2AD, CP 26, 57 rue Cuvier, F-75231 Paris Cedex 05, France »»* for allowing model training. We thank Vincent Debat for commentaries on the manuscript, and Maël Doré for help with the permutation analysis.

