## supplementary materials for "Convergence in sympatric swallowtail butterflies reveals ecological interactions as a key driver of worldwide trait diversification"

**Evaluation of the method.** Unsupervised deep learning is a powerful tool that allowed us to objectively quantify image similarity, without introducing biases due to human labelling – e.g., systematic classification such as species. However, we had to find a way to evaluate the neural network's learning. The way to evaluate SimCLR's learning is to design a downstream pretext task such as classification using only the embeddings learned from the images, allowing to assess if the information contained in the embeddings is enough to classify correctly. In our case, because of the few images per species and sex (from 1 to 5), we could not use species as labels for such a pretext task, because it led to systematic overfitting of the classifier. Instead, we found a workaround using clustering to create pseudo labels for the classification. Using a pretrained network, we clustered images into 16 groups, representing broad phenotypic groups, which we used as labels for classification. The pre-training being on the same database that our SimCLR method (ImageNet), the evaluation is not completely independent from the training. However, we do not dispose of a ground truth or enough images and so this workaround was the best way to evaluate the model. Moreover, we took care of comparing SimCLR only pre-trained with SimCLR further trained on our database, which showed improvement in performance, showing that we effectively learned information specific to our data.

| | Pre-trained-only | Pre-trained and fine-tuned | SimCLR trained |
|---|---|---|---|
| Mean accuracy | 84% | 87% | 95% |
| F1 score | 0.84 | 0.88 | 0.97 |
| Kappa score | 0.81 | 0.86 | 0.96 |

**Table. S1.** Evaluation metrics for the classification network only pretrained on ImageNet, fine tuned on our butterfly pictures, and trained with SimCLR on our butterfly pictures. Mean accuracy is the mean percentage of correct prediction over all classes. F1 score is a measure between 0 and 1 with 1 being the value for a network that perfectly predicts classes, that takes into account both precision and recall, i.e. number of true positives and false positives. Kappa score is a measure between 0 and 1, 1 being the value for a network that predicts classes perfectly, that measure the reliability while correcting for the number of classes.

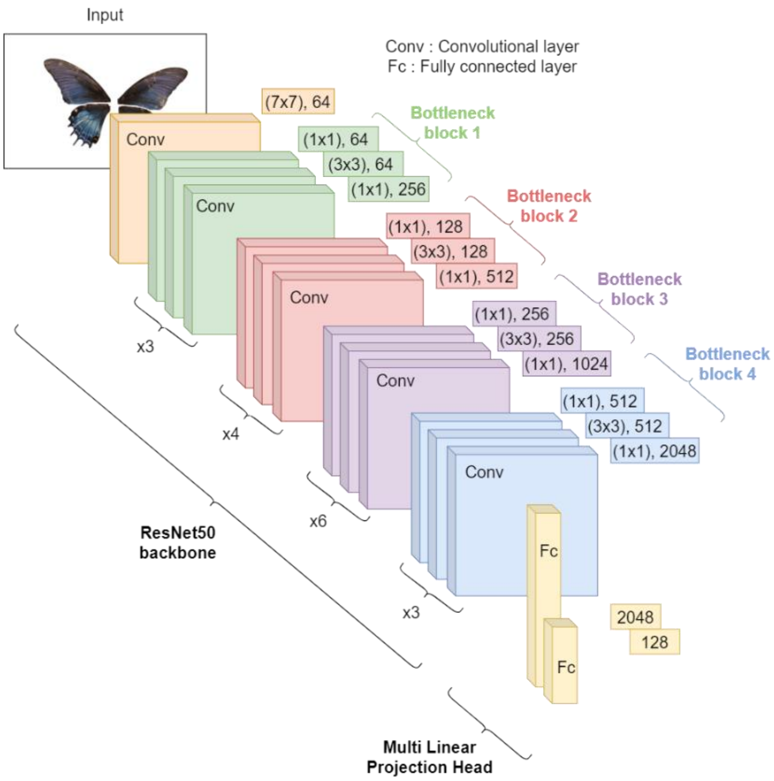


**Fig. S1.** Network architecture made of a Resnet50 convolutional neural network backbone and a multilinear projection head.

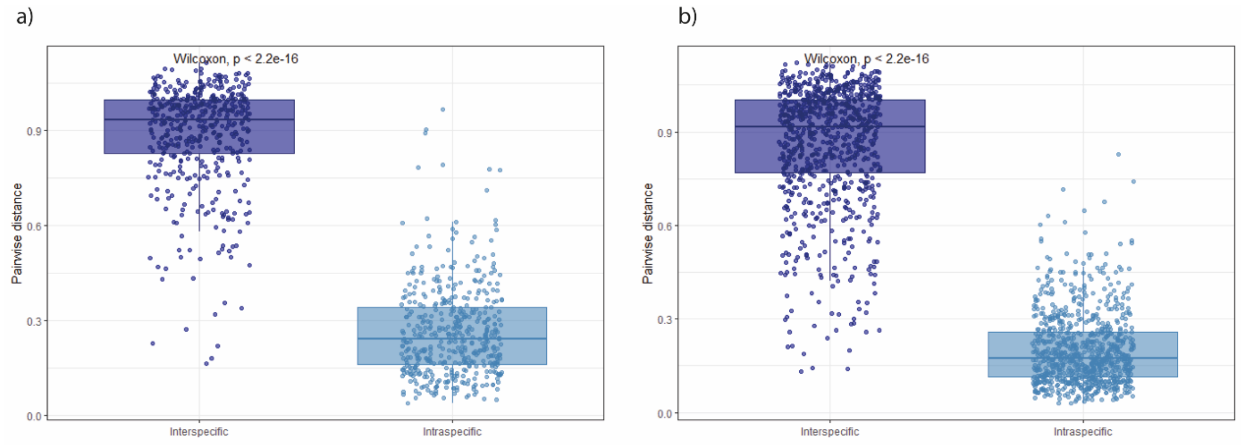


**Figure S2.** Pairwise phenotypic Euclidean distances distribution computed from the 20-dimensional morpho space coordinates, for interspecific pairs and intraspecific pairs separately for a) males and b) females. The same test was repeated for every genus separately, and among the 30 represented genus the interspecific distances were majorly higher than intraspecific distances (male dorsal sides: 23, 2 genera non-significant, 6 genera with only one species, females dorsal side: 17, 3 genus non-significant, 10 genera with only one species).

**Impact of shape on phenotypic space.** To investigate further and quantitatively the impact of wing shape on the final phenotypic space, we generated a dataset constituted by the masks of the wings for each image, filled with random colors. We then trained the method in the exact same way using this dataset, and compared the phenotypic spaces obtained. The goal was to keep the wing shape but randomize color, allowing us to separate the impact of shape from the impact of color. We computed several multivariate correlation measures between the coordinates from the two different trainings using the MatrixCorrelations R package, ranging between -1 and 1. The correlation measures showed little correlation – the maximum value being 0.27 - between the two phenotypic spaces, meaning that the wing shape did not impact much the distance measures.

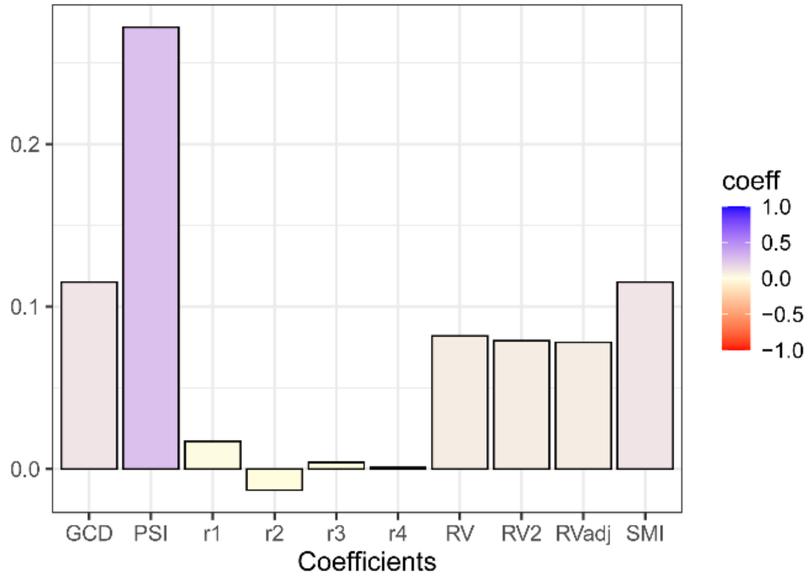


**Figure S3.** Various matrix similarity coefficients between the actual phenotypic space and the phenotypic space learned only using wing shape and randomizing colors. PSI : Procruste similarity between the two multidimensional datasets. Other statistics are listed in the MatrixCorrelation package.

**Geographical patterns of diversity and phenotypic convergence and divergence.** Phylogenetic and morphological diversity indices were computed using the EcoPhyloMapper package in R.

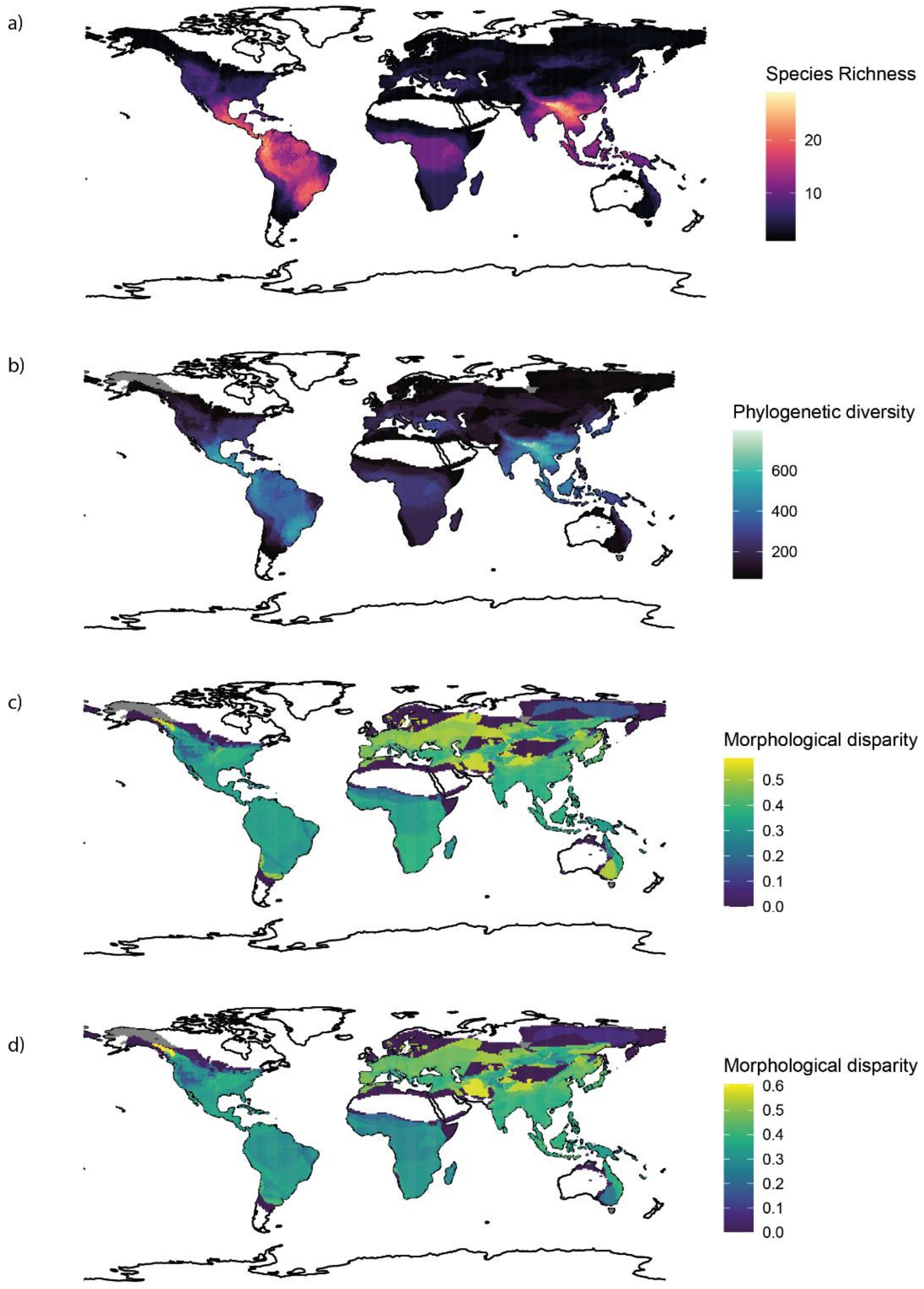


**Figure S5**: Geographical mapping of species richness (a), phylogenetic diversity (b), male phenotypic disparity (c), and female phenotypic disparity (d).

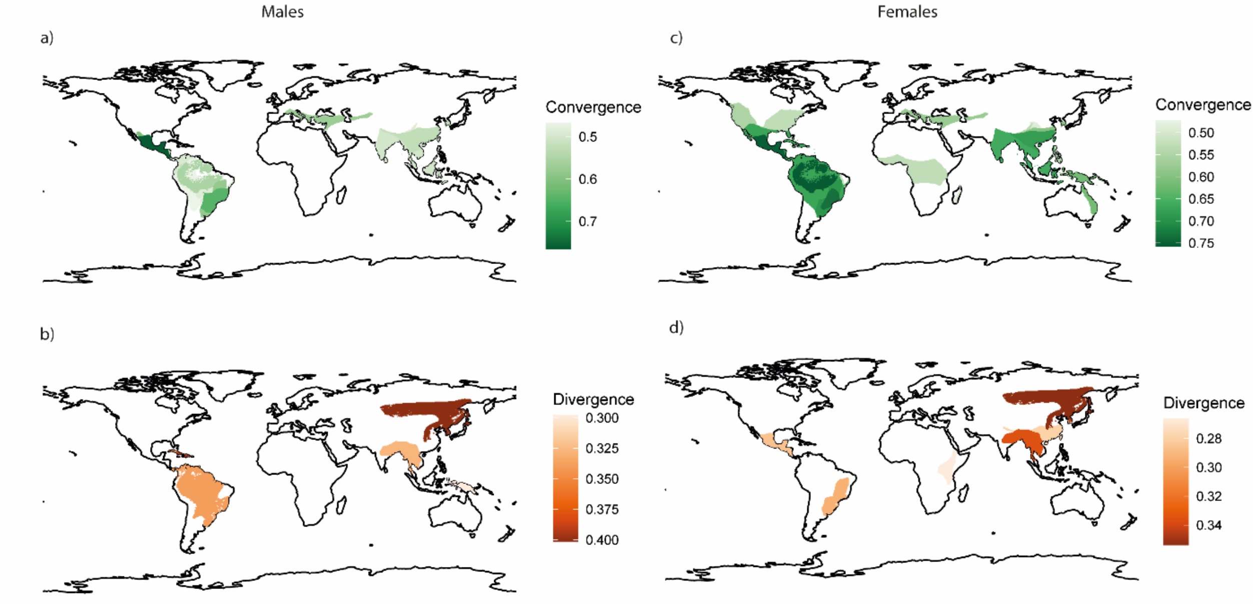


**Figure 1:** Geographical mapping of the detected convergence and divergence. Convergent and divergent areas are the intersection between range areas of the pair detected convergent or divergent. The color bar represents the maximum convergence and divergence strength. a) Mapping of male maximum convergence strength b) Mapping of male maximum divergence strength c) Mapping of female maximum convergence strength d) Mapping of female maximum divergence strength
